# Transcription Factor Co-Expression Mediates Lineage Priming for Embryonic and Extra-Embryonic Differentiation

**DOI:** 10.1101/2023.03.27.534339

**Authors:** Alba Redó-Riveiro, Jasmina Al-Mousawi, Madeleine Linneberg-Agerholm, Martin Proks, Marta Perera, Joshua M. Brickman

## Abstract

In early mammalian development, cleavage stage blastomeres and cells of the inner cell mass (ICM) of the blastocyst co-express embryonic and extra-embryonic transcriptional determinants. Using a double protein-based reporter we identify embryonic stem cells (ESC) that co-express the extra-embryonic factor GATA6 alongside the embryonic factor SOX2 in specific conditions. Based on single cell transcriptomics we find these population resemble unsegregated ICM, exhibiting enhanced differentiation potential for endoderm while maintaining epiblast competence and suggesting they represent an ideal model to determine how GATA6 and SOX2 influence each other’s DNA binding. To relate this binding to future fate, we describe a complete enhancer set in both ESCs and naïve extraembryonic endoderm stem cells and ask whether SOX2 and GATA6 recognize these elements in ICM-like ESC sub-population. Both factors support cooperative recognition in these lineages, with GATA6 bound alongside SOX2 on a fraction of pluripotency enhancers and SOX2 alongside GATA6 more extensively on endoderm enhancers. Our findings suggest that cooperative binding between these antagonistic factors both supports self-renewal and prepares progenitor cells for later differentiation.

## Introduction

How do progenitor cells sit at the cusp of two lineages, remaining stable as cell types, but simultaneously prepared for differentiation towards multiple fates? In the early mammalian embryo, the progenitors of the embryonic epiblast and extra-embryonic primitive endoderm (PrE) stably express antagonistic transcription factors (TFs) that will eventually drive epiblast and PrE lineage specification. Instead of undergoing spontaneous differentiation to both lineages, these cells exist stably across several cell cycles *in vivo* (Dietrich and Hiiragi, 2007). Here, we ask the question of how these cells express antagonistic factors and what function this might have in development and differentiation, focusing on how endoderm and epiblast enhancers become primed in different *in vitro* conditions.

Naïve embryonic stem cells (ESCs) are derived from the inner cell mass (ICM) of the mammalian blastocyst and are considered pluripotent, defined by their capacity to self-renew and to generate all the lineages of the future embryo, but not the extra-embryonic lineages (Morgani et al., 2017; Nichols and Smith, 2011). ESCs can be cultured in a range of conditions, including several defined media that support slightly different sub-populations along the spectrum of lineage specification. Culture in serum supplemented with the cytokine leukemia inhibitor factor (LIF) (serum/LIF) or defined basal media supplemented with activin, a Gsk3 inhibitor (CHIR99021) and LIF (NACL) (Anderson et al., 2017) produce cells primed towards both an epiblast and endoderm identity. By contrast, cells cultured with a MEK inhibitor (PD0325901), CHIR99021 and LIF (2iLIF) (Ying et al., 2008) homogeneously express markers related to epiblast identity, as well as containing a unique sub-population that co-express both epiblast protein and extra-embryonic RNA (Morgani et al., 2013). These cells have been reported to possess priming towards the extra-embryonic lineages by trapping ESCs with features of experimental totipotency or enhanced potency (Riveiro and Brickman, 2020). Another subpopulation that has been shown to display experimental totipotency are the 2-cell-like cells (2CLCs), a rare subpopulation that arises spontaneously in ESC culture and expresses representative factors known to be expressed in the 2-cell (2C) stage embryo (Genet and Torres-Padilla, 2020).

In this paper, we explore how co-expression of epiblast and PrE factors influences differentiation and TF occupancy. We identify spontaneous ESCs in culture that co-express SOX2 and GATA6, where these two factors govern an early ICM-like state in culture. Single cell RNA-sequencing (scRNA-seq) revealed that cells grown in Knock Out Serum Replacement media (KOSR) promote a sub-population that resembles the unsegregated ICM *in vivo*. We compile multiple sets of enhancers that characterize epiblast and PrE states *in vitro*, by using a combination of different defined ESC culture conditions (2iLIF, NACL and KOSR), as well as conditions for naïve extra-embryonic endoderm (nEnd). In ICM-like cells grown in KOSR, SOX2 is recruited to a subset of PrE-enhancers. The opposite is also observed to a lesser extent for the binding of GATA6 at pluripotency enhancers. In the subpopulation of ICM-like cells in KOSR, we observed an increase in the number of loci where SOX2 and GATA6 are co-bound. These findings suggest that cooperative binding of SOX2 and GATA6 prime enhancer states, potentially setting up competence for differentiation.

## Results

### KOSR promotes early ICM-like cells in culture

While we had previously described the co-expression of epiblast/ICM TFs with extra-embryonic RNA (Morgani et al., 2013), the co-expression of lineage opposing TFs in ESC culture is rare. To identify conditions that support a co-expressing sub-population, we generated a proteinbased double reporter ESC line with the endogenous epiblast/pluripotency factor SOX2 fused to GFP and the PrE TF GATA6 fused to mCherry (Fig. 1A), to generate SOX2-GFP/GATA6-mCherry (SGGC) ESCs (Fig. S1A-C). We confirmed that SOX2-GFP was expressed when SGGC ESCs were cultured under naïve conditions in 2iLIF and that GATA6-mCherry was expressed following differentiation towards PrE (Anderson et al., 2017) (Fig. S1D-E). We then explored whether a GATA6/SOX2 double positive (DP) population could be trapped in a variety of different culture conditions, including 2iLIF, NACL, KOSR. We also tested two culture conditions reported to produce ESCs with enhanced potency; expanded potential stem cell media (EPSCM)(Yang et al., 2017a) and extended pluripotent stem cell media containing LIF, CHIR99021, DiM and MiH (LCDM) (Yang et al., 2017b). Cells cultured in EPSCM and KOSR contained a modest fraction of DP cells (1-5%), while none of the other culture conditions supported this population at a robust level (Fig. 1B). We confirmed these findings by immunostaining (Fig. 1C), indicating that both EPSCM and KOSR cultures can support a small stable DP sub-population.

**Fig. 1.**
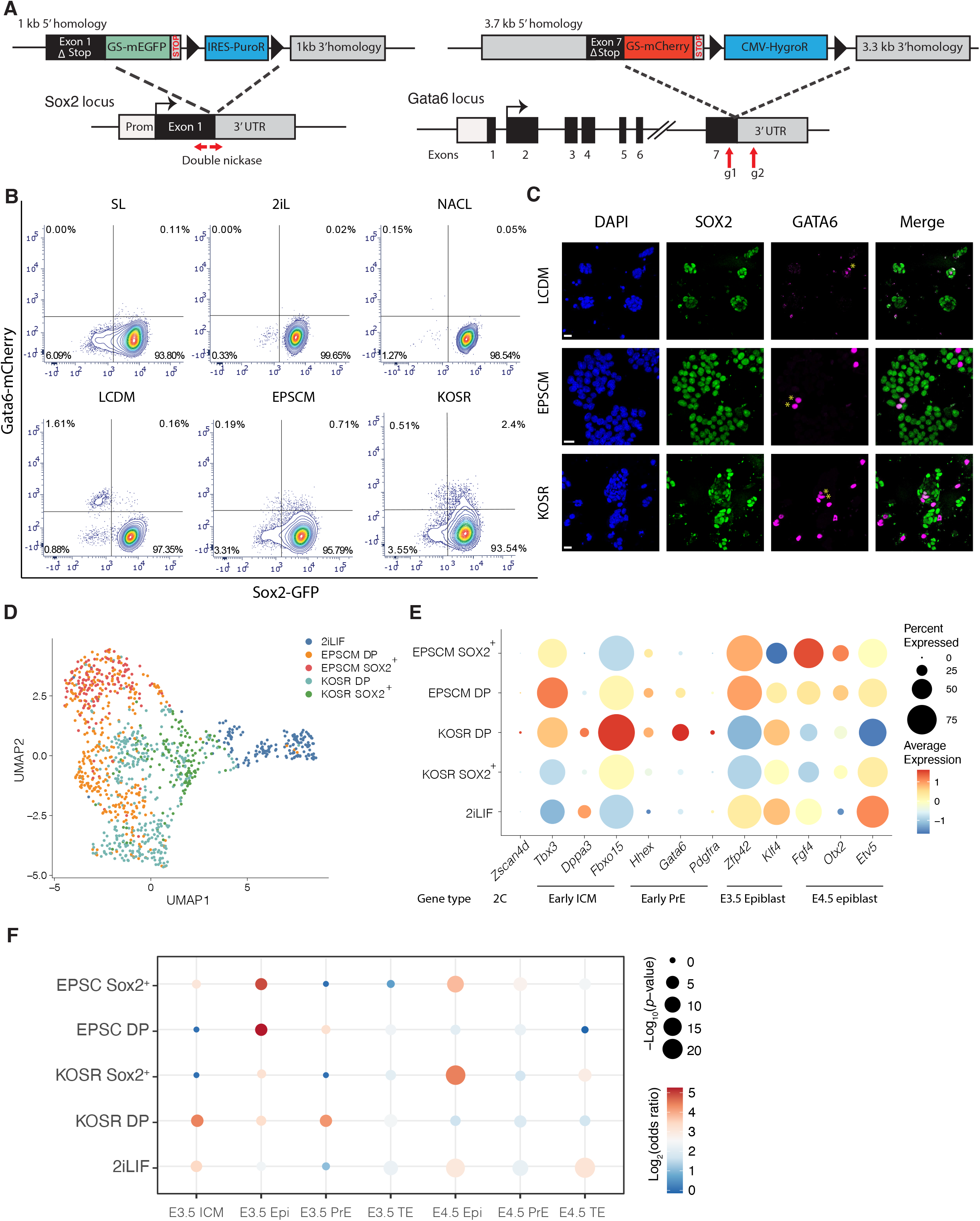
**A.** Schematic drawing of the SGG6 double reporter. **B.** Flow cytometry contour plots of the SGGC cell lines in different media conditions. **C.** Immunofluorescence images of SOX2 and GATA6 in different conditions. Scale bar = 30μm, yellow stars indicate co-expression. **D.** UMAP dimensional embedding of the scRNA-seq dataset showing the different cell populations sequenced. **E.** Dot plot graph showing the log normalized average expression of selected genes across different media conditions. The average expression is marked by the color scale and the percent of expression by the size of the circle. **F.** Gene overlap analysis of our sorted populations against differentially expressed genes from *in vivo* scRNA-seq of preimplantation blastocysts (Nowotschin et. al 2019).

To determine the nature of DP cells in EPSCM and KOSR and what their equivalent *in vivo* cell state might be, we performed scRNA-seq of SGGC cells in KOSR and EPSCM using MARS-seq2 (Keren-Shaul et al., 2019). Cells were cultured in KOSR, EPSCM or 2iLIF for at least 4 passages. DP and SOX2-GFP single positive (SOX2^+^) cells were isolated by fluorescence activated cell sorting (FACS) alongside 2iLIF control cells. After pre-processing and quality filtering, our dataset comprised 1,110 cells and 21,257 genes. Using principal component analysis (PCA), the separation of the datasets for PC1 is driven by positive expression of 2C genes (*Zscan4, Dux, Tcstv3*) while the PC2 is driven by all three culture conditions (Fig. S2A). To visualize the data, we use Uniform Manifold Approximation and Projection (UMAP) followed by unsupervised clustering. A total of 8 clusters were identified, which upon closer inspection correspond to the distinct sorted populations from the different media conditions (Fig. 1D, Fig. S2B). KOSR and EPSCM SOX2^+^ cells cluster independently from each other, whereas KOSR and EPSCM DP cells initially mix with their respective SOX2^+^ population before converging into a large DP cluster, while 2iLIF ESCs cluster separately as a branch from KOSR SOX2^+^ cells (Fig. 1D). Additionally, we observe KOSR DP cells falling into 2 clusters (cluster 1 and 2 in Fig. S2B). Cells in cluster 1 have higher levels of *Gata6*, while maintaining pluripotency marker expression (Fig S2D). Cluster 2 comprises of DP cells that have started to downregulate PrE markers (Fig S2D).

Comparing expression of candidate markers, we observe significant upregulation of both early ICM/pluripotency (*Tbx3, Dppa3, Fbxo15*) and endoderm (*Gata6*, *Pdgfra, Hhex*) genes specifically in KOSR DP cells compared to EPSCM DP cells (Fig. 1E). This suggests that the KOSR culture media is able to better capture a cell population co-expressing antagonistic lineage-specific markers of epiblast and PrE. Moreover, genes that are upregulated following epiblast specification, but before implantation, such as *Fgf4, Otx2* or *Etv5* are reduced in KOSR DP cells. However, in EPSCM DP these factors are robustly expressed (Fig. 1E).

When performing gene ontology (GO) analysis of KOSR DP compared to other populations, we observed an enrichment of terms related to metabolism. KOSR DP cells are significantly enriched for processes such as cellular response to hypoxia and mitochondrial activity, while EPSCM DP cells show enrichment for pathways related to glycolysis and pyruvate metabolic processes (Fig. S2F). KOSR DP in comparison to KOSR SOX2^+^ also show enrichment of genes related to oxidative phosphorylation (e.g. *Cox5a, Cox6c*) and lipid metabolism (e.g. *Cpt1a, Slc25a20*), as well as regulation of cell death processes and p53 activity (Fig. S2G, S3A-C). Oxidative phosphorylation and lipid metabolism are characteristic of the pre-implantation embryo, which utilizes these two as an initial source of energy before shifting to a glycolytic metabolism in the later epiblast, peaking at implantation (E4.5-E5.0) (Leese, 2012).

To assess which stages of embryo development our *in vitro* populations best represent, we compared gene expression changes in our dataset to those in the pre-implantation embryo (Nowotschin et al., 2019) by performing gene overlap analysis (Fig. 1F, S2I). Based on this, we found that the closest *in vivo* counterparts to KOSR DP cells are the E3.5 ICM and E3.5 PrE, while KOSR SOX2^+^ cells were a better fit for the E4.5 epiblast. Moreover, cluster 1 of the KOSR DP population, which express both *Gata6* and *Zfp42* (Fig. S2D), aligns to ICM, as well as both E3.5 PrE and epiblast (Fig. S2I), supporting the idea that these cells represent a dynamic ICM population, with cluster 2 of the KOSR DP representing the entrance or exit from this state. Additionally, EPSCM DP and EPSCM SOX2^+^ cells both overlapped most with E3.5 epiblast. Taken together, this analysis suggests that ESC culture in KOSR best traps a double positive SOX2-GATA6 expressing population with ICM-like characteristics, and that of the tested conditions, KOSR is the best candidate for exploring the molecular events that underlie endoderm and epiblast priming *in vivo*.

### KOSR DP cells are dynamic and primed for PrE differentiation

To study the behavior of the DP population in KOSR, we performed live imaging of steady state KOSR culture for 72 hours, a time period that is sufficient to observe DP cells arise and revert to single SOX2^+^ or single GATA6 positive (GATA6^+^) cells. Based on lineage tracking of individual cells and their descendants, we found that when DP cells arise, they maintain expression of both GATA6-mCherry and SOX2-GFP for around 2 cell cycles (~36h). Figure 2B shows the frequencies that DP cells divide into distinct daughter cells, with 57% of them maintaining their phenotype, 28% converting to SOX2^+^ and 15% into GATA6^+^. The division time of SOX2^+^ cells is fastest, the GATA6^+^ cells is slowest, with the DP population as an intermediate between the two, suggesting these media conditions are optimal for epiblast expansion. Consistent with this, the relative number of cells undergoing cell death is slightly lower in the SOX2^+^ cells (23% in DP vs 16% in SOX2^+^) (Fig. 2A, C, Video S1-2).

**Fig. 2.**
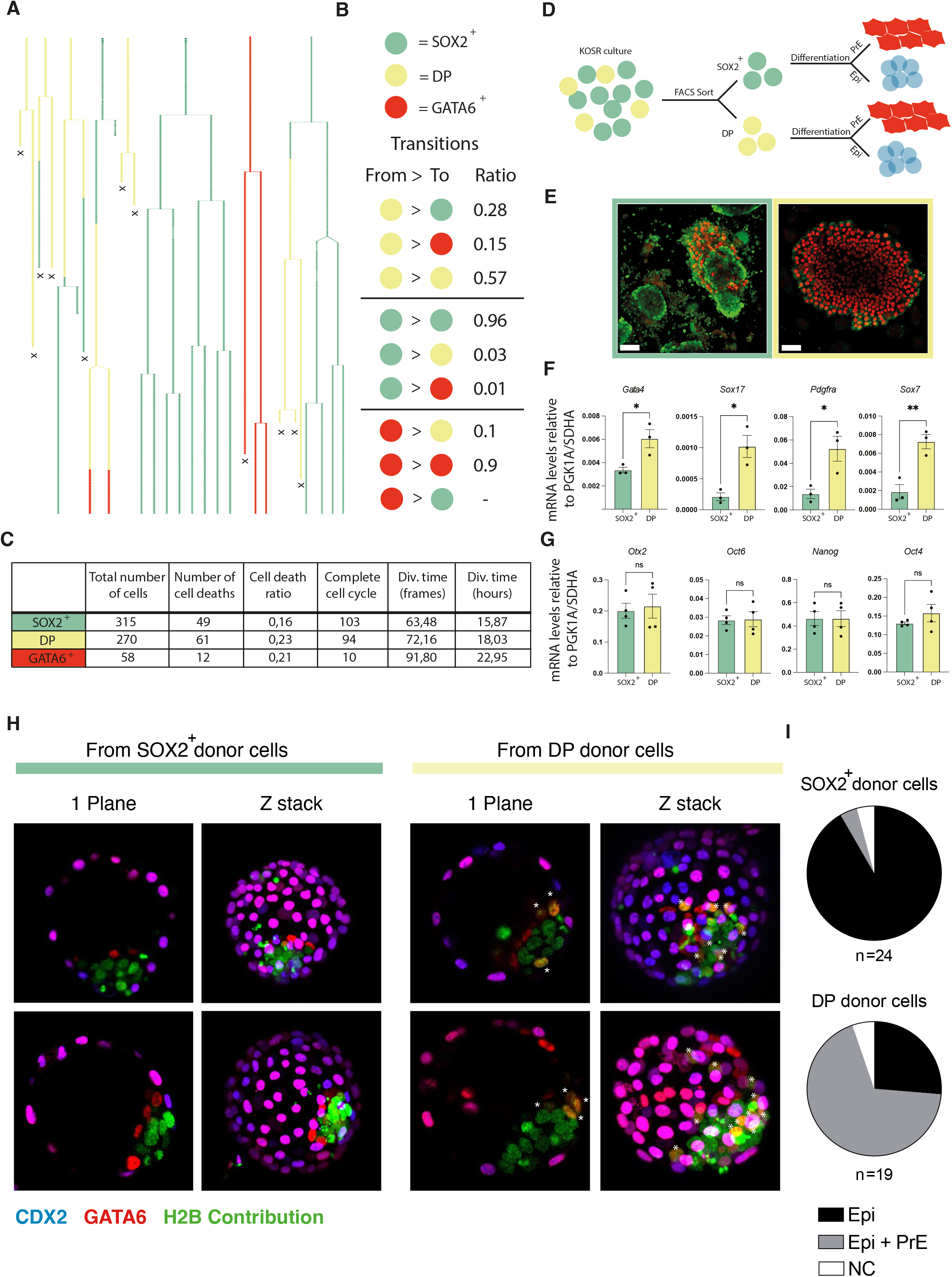
**A.** Representative dendographs examples of 8 tracked cells with its descendants (from video 22 and 24 in Suppl. Videos) for a total of 63 individual tracks of 72h. The x marks cell death. **B.** Ratios of transitions from one cell state to another. **C.** Table with quantification of the timelapse data. **D.** Representative drawing of the differentiations performed after sorting by FACS. **E.** Representative images of the sorted SOX2^+^ (green frame) and DP (yellow frame) after 5 days in primitive endoderm differentiation media. Scale bar= 50μm. **F.** Relative mRNA levels of different PrE genes of the sorted SOX2^+^ and DP after 5 days of PrE differentiation. **G.** Relative mRNA levels of different Epi genes of the sorted SOX2^+^ and DP after 3 days of Epi-like differentiation. Mean ± standard error of the mean (SEM). ns = non-significant * = p-value < 0.05 ** = p-value <0.01. **H.** Immunofluorescence images showing 1 plane or a maximum projection of E4.5 blastocysts that showed SOX2^+^ or DP donor cells contribution in morula aggregation assay. * Shows contributing donor cells expressing GATA6 **I.** Quantification of the morula aggregation assay.

Since DP cells have an overall longer cell cycle, we determined the distribution of cell cycle stages in the different population in the scRNA-seq dataset. We observed that the DP populations of both EPSCM and KOSR have longer G1 and G2 phases (Fig. S3A). As cells primed for PrE differentiation have been shown to stay for longer in G1 (Coronado et al., 2013; Perera et al., 2022), this could explain why DP cells possess a longer G1 phase. An increase in the G2 phase may be linked to an increase in cell death in the DP cells, as G2 arrest normally proceeds apoptosis. (Pietenpol and Stewart, 2002).

Given the dynamic nature of the DP population, we reasoned that spontaneously occurring ICM-like cells could represent an intermediate in PrE differentiation that are also capable of giving rise to epiblast. Therefore, ICM-like DP cells should exhibit an enhanced capacity or bias to undergo PrE differentiation, relative to single SOX2^+^ ESCs, but this should not be at the expense of a reduction in the efficiency with which they differentiate to epiblast. To test this hypothesis, we assessed the relative efficiency of KOSR DP and SOX2^+^ cells to differentiate into PrE (Anderson et al., 2017) and to mature epiblast, or Epi-like cells (Hayashi et al., 2011) (Fig. 2D). Figure 2E shows that PrE differentiation from sorted DP cells rapidly produce robust PrE monolayers within 5 days, whereas SOX2^+^ cells only partially differentiate, an observation that is supported by RT-qPCR for PrE markers (Fig. 2F). In contrast, the efficiency of epiblast differentiation of these two populations is not significantly different (Fig. 2G). We also observe a small, but inconsistent capacity of the DP cells to upregulate TE markers upon differentiation (Tanaka et al., 1998) (Fig. S4B-C). Finally, we also found that individual SOX2^+^ and DP cells have a similar capacity to support the expansion of undifferentiated colonies in clonal assays (Fig. S4B, D-E). Since DP cells can readily differentiate to Epi-like and PrE, mimicking ICM identity, we performed morula aggregation with H2B-mir670 tagged SOX2^+^ and DP cells and analyzed their contribution to host blastocysts. As expected, SOX2^+^ cells contributed extensively to the epiblast (22/24 embryos), while the DP cells were found predominantly in both epiblast and PrE lineages (13/19 embryos) (Fig. 2H-I). Taken together, these observations support the existence of a transient ICM-like state in KOSR culture.

### Co-expression of SOX2 and GATA6 induces changes to canonical binding

To determine how these two antagonistic TFs could be co-expressed in KOSR and whether they might reciprocally influence each other’s binding or activity, we sought to identify the extent to which they recognize the same target in different populations. While considerable effort has been invested into understanding the pluripotency or epiblast network (Li and Belmonte, 2017), by comparison, little has been invested into PrE. To provide a framework by which we can understand the nature of this lineage bifurcation, we first established the enhancer network in differentiated PrE cells and compare it to the epiblast, as recapitulated in different pluripotent culture conditions. We assessed the enhancer network in nEnd stem cells, and compared them to two distinct states of pluripotency: 2i/LIF and NACL. NACL is a defined pluripotent culture system where cells exhibit the same heterogeneity as conventional serum containing media. In addition, NACL media is composed of the same set of cytokines as used for nEnd culture, but differs only in its base media (N2B27) (Anderson et al., 2017). We used Cleavage Under Targets & Release Using Nuclease (CUT&RUN) (Skene and Henikoff, 2017) to assess the status of regulatory regions that are co-occupied by histone modifications known to indicate enhancer activity, H3K27ac and H3K4me1 (Calo and Wysocka, 2013) (Fig.3A-E). Based on the combination of these marks in two clonal cell lines in both pluripotent conditions, we identified 6849 active pluripotency enhancers (Fig.3A, B, C). In nEnd, we found 4957 active PrE enhancers, with 1434 of these being shared by both lineages, referred to as “common enhancers” (Fig. 3B, D, E). Motif analysis indicated that these regulatory regions are cell type specific as we identified specific groups of motifs enriched in each enhancer set. We observed that the PrE subset was highly enriched for GATA motifs, while the pluripotency subset enriched for the OCT, NANOG, ESRRB and KLF motifs. The common enhancer subset feature a strong KLF signature, consistant with its expression in both lineages (Morgani and Brickman, 2015; Nowotschin et al., 2019) (Fig. 3F). GO analysis for Biological Processes for the closest gene to the different subset of enhancers supports that “common enhancers” represent general cellular processes, implying significant numbers of housekeeping genes (Fig. 3H). The “pluripotency enhancers” subset includes both LIF response and embryo development terms (Fig. 3G) and in the “PrE enhancers” have an abundance of terms related membranes and adhesion in accordance with nEnd cells undergoing morphological changes (Fig. 3I).

**Fig. 3.**
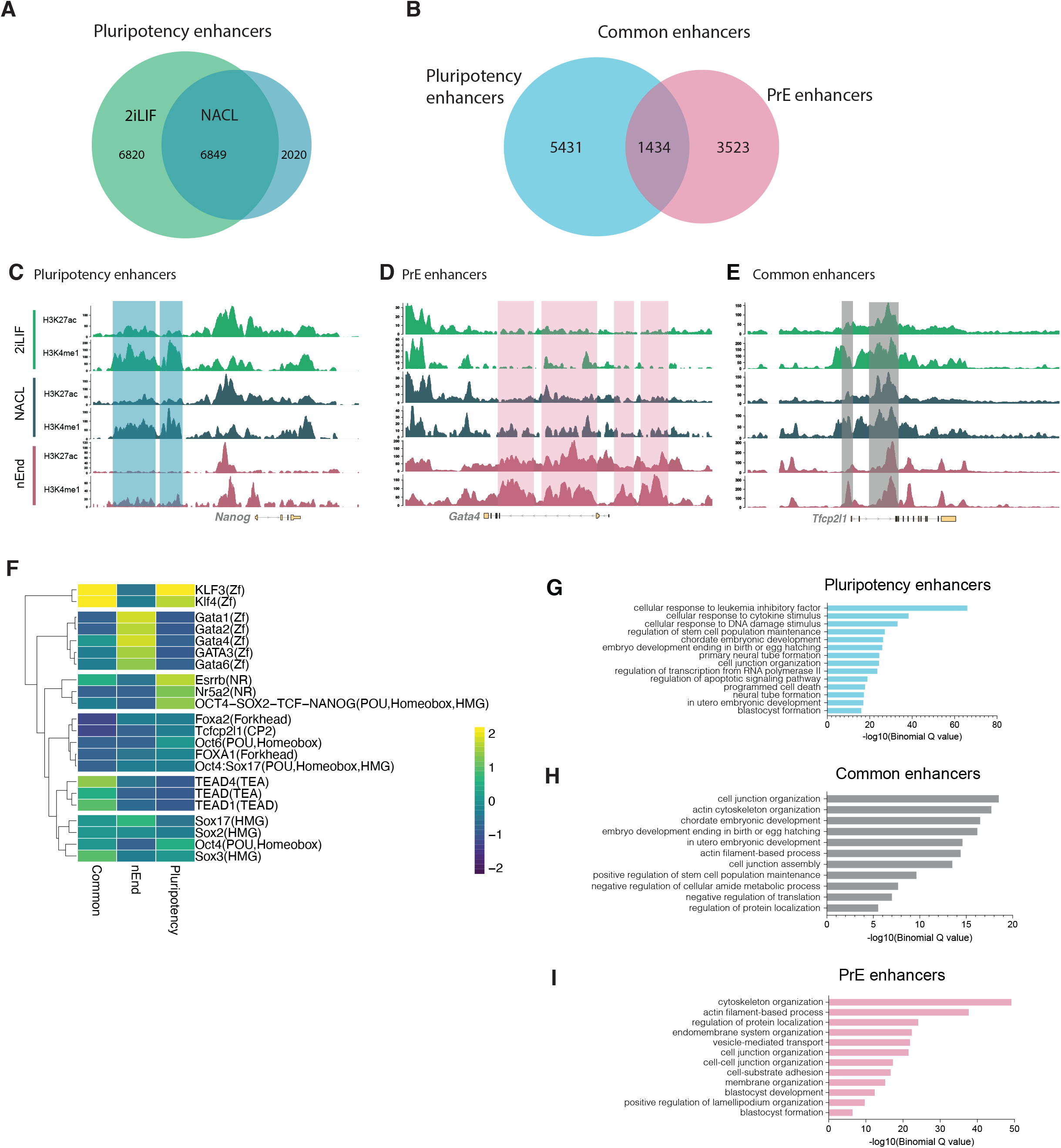
**A.** Euler diagram defining an intersect of pluripotency enhancers: enhancers active in both 2iLIF and NACL. **B.** Euler diagram comparing pluripotency enhancers (intersect from A) with PrE enhancers. **C-E.** Genome browser tracks (Gviz package) of H3K27ac and H3K4me1 across conditions 2iLIF, NACL and nEnd of loci *Nanog* (C), *Gata4* (D), and *Tfcp2l1(E*) with examples of pluripotency enhancers (blue), PrE enhancers (pink) and common enhancers (grey) defined in B. **F.** Heatmap of selected motifs across enhancer subsets defined in B, with a cutoff of p-value < 0.0005. Scale shows −log10(p-value). **G-I:** GO terms of the biological processes of pluripotency enhancers (G), common enhancers (H) and PrE enhancers (I).

Having established the enhancer networks in both lineages, we assessed TF binding at both enhancer sets in both the final cell states (ESC and nEnd), and in the DP and in SOX2^+^ populations cultured in KOSR. We used CUT&RUN to probe the binding of both factors in all the conditions in which they were expressed; GATA6 in nEnd and DP populations and SOX2 in NACL, 2iLIF and in the SOX2^+^ and DP cells in KOSR. In order to determine if SOX2 and GATA6 binding shifts globally, we compared SOX2 and GATA6 binding on pluripotency, nEnd and common enhancer sets (Fig. 4A). While we observed a decrease in SOX2 binding to pluripotency enhancers in DP cells, we observed a significant recruitment of SOX2 to PrE enhancers alongside GATA6, suggesting that SOX2 binding can move towards a potential GATA6 site. We also observed a smaller acquisition of GATA6 peaks at pluripotency enhancers, slightly greater than GATA6 binding the same elements in nEnd (Fig. 4A). We detect 1716 peaks cobound by SOX2 and GATA6 in the DP cells (Fig4. B), and 416 of these peaks (24%) sit at enhancers that we previously defined. Interestingly, 50% of the cobound peaks sit at PrE enhancers, while only 28% of these are found at pluripotency enhancers and 22% at common enhancers. Taken together, this suggests that SOX2 is recruited to sites with GATA6 occupancy, and when they sit at enhancers, this predominantly occurs at PrE enhancers. Given the PrE bias in occupancy, we analyzed the closest genes regulated *in vivo* (Boroviak et al., 2018) to these SOX2-GATA6 cobound peaks and observed a 2-fold enrichment of PrE genes over epiblast genes. Moreover, in the cobound regions, regardless of their affiliation to lineage specific genes, contain twice as many GATA6 motifs than those found for SOX2 (Table 1 and 2). Only around 10% of the closest genes to the cobound peaks are significantly upregulated when DP cells are compared to SOX2^+^ single positive cells (specifically 4,9% for epiblast genes and 11.7% for PrE genes, Table 1 & 2), suggesting that lineage priming is not occurring at the transcriptional level.

**Fig. 4.**
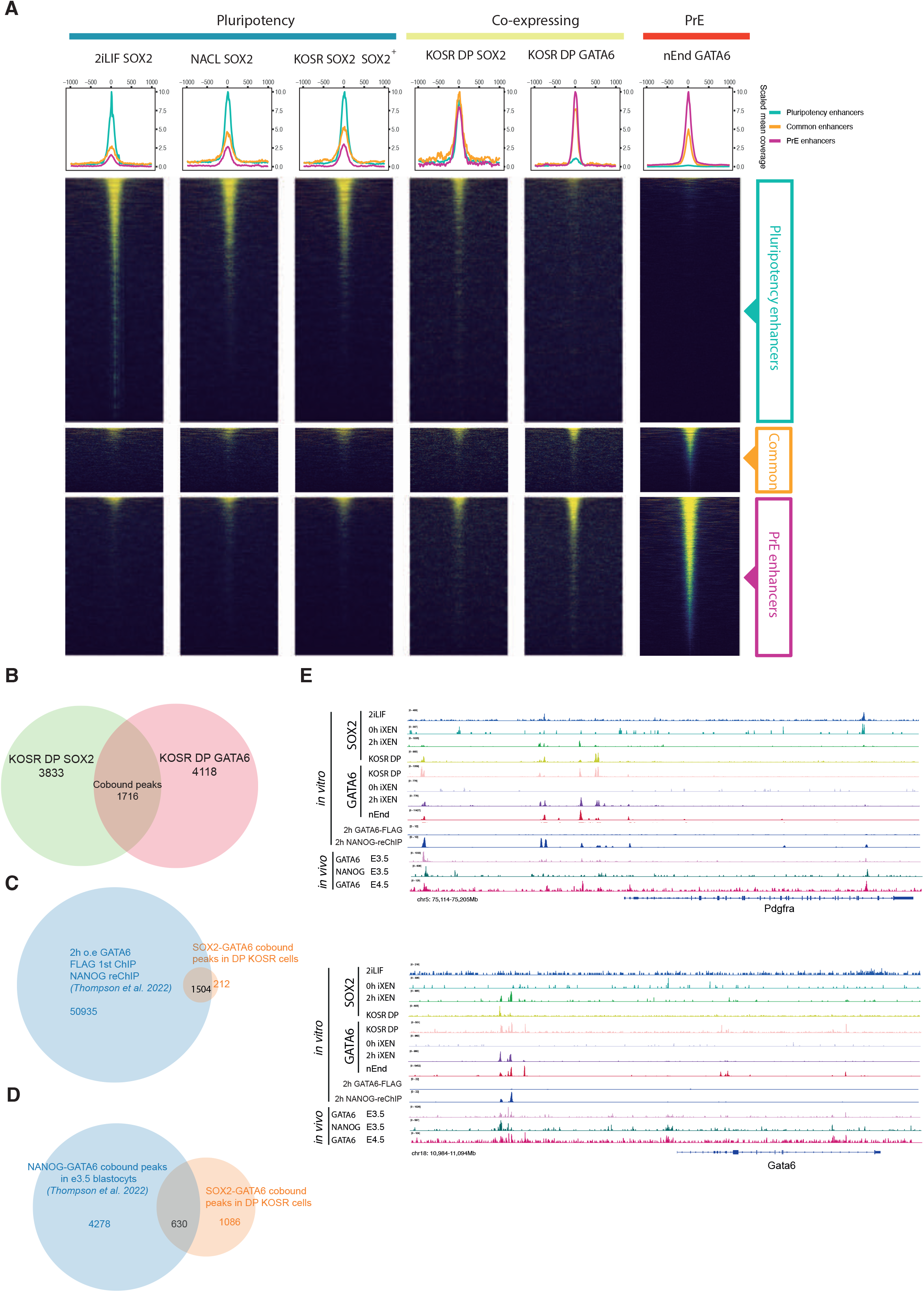
**A.** Scaled heatmaps showing SOX2 and GATA6 binding at the pluripotency, common or PrE enhancer subsets defined previously, in the conditions; 2iLIF, NACL, KOSR sorted cells and nEnd. **B.** Euler diagram of SOX2 and GATA6 peaks in KOSR DP cells. **C.** Euler diagram showing the overlap of the GATA6-FLAG ChIP with NANOG reChIP (*Thompson et al. 2022*) with our SOX2-GATA6 cobound peaks in DP cells. **D.** Euler diagram projecting the NANOG-GATA6 cobound peaks from the *in vivo* data of the E3.5 blastocysts (*Thompson et al. 2022*) with the SOX2-GATA6 cobound peaks in KOSR DP cells. **E.** Genome browser tracks (IGV 2.14.0 software) of SOX2 and GATA6 comparing data from this study (2iLIF, KOSR and nEnd tracks) with *Thompson et al. 2022* (0h and 2h iXEN induction, the GATA-FLAG ChIP with NANOG reChIP and the E3.5 and E4.5 blastocysts tracks). *Pdgfra* locus (top panel) and *Gata6* locus (bottom panel).

While our data on cobinding is not based on sequential precipitation, we took advantage of a recent study that explored a similar cobinding phenomena by sequential ChIP in response to GATA6 induction in ESCs to drive differentiation to iXEN cells (Thompson et al., 2022). We observed a significant percentage of overlap of our binding for SOX2 and GATA6 in DP cells with this study’s early timepoints for GATA6 induction (Fig 4E and Table 3). Here we found that between 70-80% of our DP peaks overlap with the 2 and 4h overexpression time points for the factor GATA6 and 50-60% for SOX2. We observe an impressive 87% overlap of our cobound peaks with the NANOG-GATA6 cobound peaks at 2h based on sequential precipitation (GATA6-FLAG-NANOG) (Fig. 4C, E), suggesting that our SOX2 and GATA6 are indeed binding at the same sites. We also observe a 37% overlap of our DP GATA6-SOX2 cobound peaks with their NANOG-GATA6 cobound peaks in the *in vivo* E3.5 blastocysts (Fig 4D). Moreover, assessment of this data also suggests that the preference of SOX2/GATA6 binding for PrE enhancers in our data may be a general property of cooperative binding between endoderm and epiblast determinants, as *in vivo* ICM GATA6-NANOG cobound peaks also appear to more often bind in our endoderm enhancers (48%) compared to their presence at at pluripotency ones (33%).

## Discussion

In this study, we found that ICM-like expression of endoderm and epiblast TFs prepare differentiation not only for their own lineage, but for the reciprocal as well. Cooperative binding interactions between SOX2 and GATA6, simultaneously prepare enhancer for activation, while maintaining these genes in a primed status that enables rapid response in lineage bifurcation. Multipotent progenitors must maintain tight control of lineage commitment to ensure correctly proportioned embryonic development. Lineage-specific inducer genes such as GATA6 and SOX2 that are co-expressed in early ICM cells (Dietrich and Hiiragi, 2007; Morgani and Brickman, 2015), could prime later differentiation based on their ability to stabilize each other’s binding and maintain cells with the ability to kick start either lineage specific gene regulatory network.

DP cells can readily differentiate towards epiblast or PrE, while their sister cells in culture (SOX2^+^ cells) appeared to be biased towards epiblast. Transcriptionally and metabolically DP cells also best approximate the E3.5 unsegregated ICM, whereas the SOX2^+^ cells resemble the E4.5 epiblast. In agreement with (Posfai et al., 2021), we observe that cells cultured in EPSCM present a more epiblast signature. Given that the ICM can sustain the co-expression of these factors over multiple cell divisions, the wiring of these cooperative interactions between cross lineage TFs are likely to be modestly stable, and here we find that *in vitro* ICM-like cells can sustain this state through cell division as well. A recent study based on GATA6 over-expression, describes an *in vitro* model for the ICM. In this instance, this ICM-state is hypothesized to exist at the early time points of GATA6 mediated reprogramming of naïve ESCs (Thompson et al., 2022). However, here we manage to capture a small population of cells that intrinsically appears able to trap this state in steady state culture without artificial manipulation of gene expression. The ability to sort this population at a steady state enabled us to correlate differentiation competence with TF cooccupancy. Further work is required to properly understand the ICM’s environment, which in turn will lead to better ways to robustly expand these cells *in vitro*. But even as a small population these cells appear to represent a genuine stem cell model that maintains embryonic extra-embryonic bipotency indefinitely.

We also defined the enhancer network in nEnd relative to naïve pluripotent ESCs (2iLIF and NACL). Unsurprisingly, the binding motifs of the pluripotency set of enhancers are enriched in genes of partaking in pluripotency network, such as OCT4, SOX2, NANOG, KLFs and ESRRB, while nEnd cells are enriched mostly in GATA motifs. Common enhancers, which have H3K27ac and H3K4me1 signal in both nEnd and naïve cultures, are enriched for KLF motifs. While we observed GATA6 and SOX2 binding to all these enhancer sets, we observe an almost identical bias in our data and that derived from the E3.5 ICM *in vivo* (*Thompson’s data*) for PrE enhancers, as well as 2 fold increase in GATA6 motifs over SOX2 motifs. This suggests that GATA6 binds its consensus sites with a relatively high affinity and actively recruits SOX2, rather than the other way round. This would appear to contrast its role in ESCs, where residence time data and *in vivo* imaging studies that suggest that SOX2 drives OCT4 binding (Chen et al., 2014; White et al., 2016). Alternatively, this might reflect that ability of SOX2 to recognize lower affinity elements, like those recognized by SOX17, only in presence of GATA6. Moreover, we observed the presence of SOX sites in the proximity of GATA in the PrE, but not the other way round. Yet, why then should the presence of GATA6 facilitate both endoderm and epiblast differentiation? Perhaps the relative levels of free SOX2 is a key determinant of epiblast differentiation, and the presence of GATA6 titrates SOX2 away from epiblast enhancers and OCT4, maintaining a threshold concentration that can be pushed toward either endoderm or epiblast.

In hematopoietic differentiation, the co-expression of antagonistic lineage specifiers is thought to maintain progenitor populations at the apex of two lineages based on a phenomenon known as multi-lineage priming. Here we find that cooperativity between GATA6 and SOX2 leads to alterations in their binding in DP cells, such that they are sitting at sites found in both lineage specific enhancer sets and presumably there is neither sufficient levels of these factors to drive differentiation or perhaps the expression of these two TFs in the absence of lineage determining signaling is insufficient to promote differentiation (Hamilton et al., 2019; Knudsen and Brickman, 2020). Is the combination of these factors cross-antagonistic at the level of direct transcriptional regulation? Based on the observation that these factors bind together to enhancers that drive transcription in both cell types, makes this seem unlikely. This suggests that multi-lineage priming may be about manipulating threshold distributions for lineage specification, exploiting antagonistic factors with known interactions and capacity of cooperativity to support cells in a precarious balance between opsoing fates.

## Data availability

scRNA-seq and CUT&RUN data that support the findings of this study have been deposited in NCBI GEO with the code GSE227889, which the reviewer can access with the token: **cpyxwseefpqlhaz** (https://www.ncbi.nlm.nih.gov/geo/query/acc.cgi?acc=GSE227889)

## Acknowledgments

We thank Paul van Dieken and Gelo de la Cruz for flow cytometry and FACS support and expertise. We thank Michaela M. Rothová, Helen Neil and Magali Michaut for technical expertise and support with the scRNA Seq and CUT&RUN sequencing. We thank Javier Martin Gonzalez and Ricardo Alonso Laguna Barraza and all the transgenics platform facility for technical assistance and providing mouse embryos. We thank Jutta Bulkescher and the reNEW Imaging Platform for training, technical expertise, support, and the use of microscopes. We thank Jose Alejandro Herrera Romero and Sarah Louise Lundegran for bioinformatics advice. Lastly, we thank the entire Brickman lab for critical discussion. This work was support by Lundbeck Foundation (R198-2015-412 and R370-2021-617), Independent Research Fund Denmark (DFF-8020-00100B), and the Danish National Research Foundation (DNRF116). ARR and MPP were supported by a Lundbeck Foundation PhD studentships (R208-2015-2872 and R286-2018-1534). The Novo Nordisk Foundation Center for Stem Cell Medicine (reNEW) is supported by a Novo Nordisk Foundation grant number NNF21CC0073729, and previously NNF17CC0027852.

## Author Contributions

ARR and JAM performed and analyzed experiments. MLA, MPP, MP analyzed experiments. JMB supervised the project. ARR and JMB designed the project and wrote the paper with input from all other authors.

## Materials and Methods

### ESC culture

ESCs were generated using E14JU ESCs from the 129/Ola background. ESC lines were maintained in serum/LIF (Canham et al., 2010), 2i/LIF (Ying et al., 2008), KOSR (Martin Gonzalez et al., 2016) EPSCM (Yang et al., 2017a) and LCDM (Yang et al., 2017b) as previously described.

### Generation of SOX2::GFP-GATA6::mCherry (SGGC) ESC lines

SOX2:GFP ESCs (Anderson et al., 2017) were used to further target with a Gata6:mCherry construct using CRISPR-Cas technology. The construct contains mCherry tagged immediately after exon 7 of the Gata6 locus, just before the STOP codon and the construct has 3000bp homology arms.

SOX2:GFP ESCs were plated into a gelatinised 6-well plates in SL and lipofected with the linearized Gata6:mCherry-Hygromycin plasmid and the CRISPR plasmid using Lipofectamine2000 (ThermoFisher 11668019) following the manufacturer’s instructions. After 16 h incubation, the cells were transferred into 10cm dishes with fresh medium supplemented with Hygromycin to select for colonies successfully integrated. We obtained 3 clones (B9, B12 and E1) that were successfully integrated. Clone B9 was generated using CRISPR guide 1 (GCTCTGGCCCTGGCC), which cuts at the last 20nt of the coding sequence of Gata6, and clones B12 and E1 were generated using CRISPR guide 2 (GCACAGAAATCACGCATCGA), which cuts 150 bp after the STOP codon (See Fig. 1A).

SGGC cells were verified by performing immunostaining for SOX2, GATA6, mCherry and GFP, Western blot, locus sequencing to screen for unwanted mutations generated by CRISPR, karyotyping and Southern blot.

### Karyotyping

SGGC ESCs were expanded until 50-60% confluency, after which they were incubated for 1h in medium containing 10 μg/ml Colcemid (Sigma Aldrich D1925). The medium was collected for separate disposal, and without a PBS washing step, 2 ml of trypsin were added. When the first cells started to detach, the trypsin was inactivated by adding medium. Pelleted cells were resuspended in 2.5 ml hypotonic solution (0.56 % (w/v) KCl and incubated at RT for exactly 6 min. 1 ml of fixative (75 % (v/v) methanol, 25 % (v/v) acetic acid) was added, followed by 30 min incubation at 4°C. After 2 washes in fixative, cells were carefully resuspended in 200μl fixative and spread onto pre-cleaned poly-L-lysine coated glass slides. Chromosomes were stained for 30 min in filtered 10% Giemsa pH 7.2 solution and imaged using a 63xOil objective on an inverted Olympus microscope.

### Southern blot

Southern Blotting was used to test SGGC clones for correct gene targeting as previously described (Southern, 2006). Gata6:mCherry construct integration was checked using both an internal (mCherry) and an external probe. DNA for the external probe was cut using AflIII and NdeI. The DNA for mCherry internal probe was cut using HindIII (Fig. S1A-B). Hybridization of the probe happened at 60°C O/N. The blots were left in the exposure cassette for 48h and developed using a high-resolution Typhoon scanner.

### Flow cytometry and FACS

Cells were washed in PBS and brought to single cell suspension using Accutase. Cells were resuspended in PBS/FCS with DAPI (1:10,000) to exclude dead cells. Cells were analyzed using a LSR Fortessa flow cytometer (BD Biosciences) and the FACSDiva (BD Biosciences) software. Plots were generated using FCS Express 6.0 (DeNovo Software). Cells were sorted by SOX2^+^ or DP populations using a BD FACS Aria III (BD FACSDiva Software version 8) with a 100 μm nozzle. The boundary between positive and negative populations was set on the basis of negative population of control ESCs. We also used parental line SOX2::GFP (SG16) to set the gates for mCherry positive cells.

### Differentiations

ESCs were cultured for four passages in KOSR prior to differentiation, isolated by FACS for DP or SOX2^+^ expression and seeded at the same cell density for both. Upon attachment, media was replaced for specific differentiation conditions.

For nEnd differentiation, we plated 6 × 10^4^ cells/cm^2^ in gelatinized plates and cultured the cells in RACL media as previously described (Anderson et al., 2017; Linneberg-Agerholm et al., 2019).

For Epi-like differentiation, we plated 20 × 10^4^ cells/cm^2^ in fibronectin-coated plates and cultured the cells in Epi-like media for 3 days (Hayashi et al., 2011).

For TSC differentiation, we plated 10 × 10^4^ cells/cm^2^ and then transferred to trophoblast stem cell medium for a total of 6 days (Tanaka et al., 1998).

### Alkaline Phosphatase Staining

ESCs were plated at clonal density and cultured for 7 days. Alkaline phosphatase staining was carried out as per manufacturer’s instructions. Colonies were imaged using a Leica M165 C microscope.

### ESC and blastocyst immunostaining

ESCs were cultured in 8-wells slides (Ibidi) and immunostaining was carried out as previously described (Canham et al., 2010). The following antibodies were used anti-SOX2 (Santa Cruz sc17320) used 1:200, anti-GATA6 (Cell Signalling Technologies, 5851) used 1:1500. Secondary antibodies used are from the Alexa fluor series (Molecular Probes, ThermoFisher), all 1:2000. Blastocysts were stained by anti-CDX2 (BioGenex MU392A-UC) used at 1:200 and anti-GATA6 (Cell Signalling Technologies, 5851) used at 1:200. Both mESCs and blastocysts were imaged using a confocal Leica SP8.

### Quantitative PCR with reverse transcription (RT-qPCR)

Total RNA was collected using the RNeasy Mini Kit (Qiagen). 1μg of total RNA was used for first strand synthesis using SuperScript III reverse transcriptase according to the manufacturer’s instructions. cDNA corresponding to 10ng total RNA was used for RT–qPCR analysis using the Roche LC480 and target amplification was detected with the Universal Probe Library system.

### scRNA-seq

Cells were sorted by FACS using a BD FACS Aria III with a 100μm nozzle and 20psi sheath pressure. The boundary between positive and negative populations were set based on the negative population of control ESCs. A forward scatter (FSC) and side scatter (SSC) was used to define a homogeneous population, FSC-H/FSC-W gates were used to exclude doublets, and dead cells were excluded based on DAPI staining. Cells were sorted directly into 384-well plates containing lysis buffer which includes the first RT primer and RNase inhibitor, immediately frozen and later processed by the MARS-seq2 protocol (Keren-Shaul et al., 2019). scRNA-seq libraries were sequenced using Illumina NextSeq500 at a median sequencing depth of 225,000 reads per single cell. Pre-processing was done using nfcore/marsseq pipeline (Proks 2023) with following command: nextflow run . -c dangpufl01.config --genomes_base ./references --input ./Alba_SB2/design.csv

### Downstream analysis of single-cell dataset

Both *in vivo* (Nowotschin et al., 2019) and *in vitro* datasets were processed using Seurat (v4.1.0). Based on the quality control for the *in vitro* dataset, we filtered cells using following cutoffs: 1,300 < # genes < 7,500 and 500 < # UMI < 40,000 and mitochondrial content > 20%. Empty control wells labeled as Zero and ERCC-genes were also discarded. The filtered dataset containing 1,110 cells were further normalized using ‘sctransform’ workflow followed by identifying 3,000 highly variable genes with default settings. UMAP was computed for first 20 PCAs, followed by Louvain unsupervised clustering (‘FindClusters’), which estimated 8 clusters with a set resolution of 1. Top differentially expressed genes were identified using Wilcoxon-rank sum test with following cutoffs (log2foldchange > 1 and p-value < 0.05). The full analysis can be found at https://github.com/brickmanlab/riveiro-et-al-2022/. Gene overlap plots were created with “Shen L, Sinai ISoMaM (2021). *GeneOverlap: Test and visualize gene overlaps*. R package version 1.30.0, http://shenlab-sinai.github.io/shenlab-sinai/.”

### Time-lapse imaging and cell tracking

SGGCH2B ESC lines were cultured in KOSR media, on 8-well slides (ibidi) and imaged every 15 minutes across 72h. mCherry, GFP and CY5 fluorescent light channels were used in 5%CO2, 20% O2 at 37°C under a Deltavision Widefield Screening microscope. ESCs were seeded at 5000 cells/cm^2^ 24h before the beginning of the time lapse in KOSR. We performed manual cell tracking using Imaris v9.5 (Bitplane). Nuclei were segmented using the H2B marker tagged with far red. We measured the SOX2-GFP and GATA6-mCherry fluorescence intensities of a circular area of 50 μm diameter inside the segmented nuclei. For each area measured, we took the median fluorescence intensity as the measure for that given data point. Intensity measurements were linked to its time point and lineage, allowing us to infer the division time for each cell that was tracked, as well as the expression level of both SOX2 and GATA6 in each time point. Only cells with completed cell cycle information were used for calculate the transition analysis. A total of 63 individual tracks of 72h have been tracked.

### CUT&RUN

KOSR, nEnd, 2iLIF or NACL cells were grown in their respective media for at least 4 passages. KOSR and nEnd cells were sorted by FACS using a BD FACS Aria III. From KOSR we sorted SOX2^+^ and DP cells. From nEnd we sorted GATA6^+^ cells. A minimum of 100,000 cells were sorted to proceed with the CUT&RUN protocol. CUT&RUN was performed using an in house purified MNase and following the published protocol (Janssens and Henikoff, 2019). Library preparation was performed following this published protocol (Liu, 2019). The quality of the CUT&RUN libraries was assessed using an Agilent Fragment Analyzer and they were quantified using Qubit. Sequencing was performed on an Illumina NextSeq 550, up to 48 uniquely barcoded samples were pooled on a High Output flow cell. Samples are sequenced using paired end sequencing. Antibodies used: Sox2-Active Motif 39843, Gata6-Cell Signaling 5851, H3K27ac Abcam-ab4729, H3K4me1 Abcam-ab8895. All used at 1:100 concentration.

### Downstream analysis of CUT&RUN

The software used for pre-processing was Cutadapt (Martin, 2011), Bowtie 2 (Langmead and Salzberg, 2012), Picard2, SAMtools (H et al., 2009), BEDtools (Ar and Im, 2010) and DeepTools (Ramírez et al., 2016). All downstream data analysis was performed using BEDtools, SAMtools, DeepTools, Fluff (Georgiou and van Heeringen, 2016), Integratative Genome Viewer (Robinson et al., 2011) SEACR (Meers et al., 2019), HOMER (Heinz et al., 2010) and RStudio (RStudio, 2016). Peaks were called using SEACR, parameters; ‘relaxed’ for transcription factors and ‘Stringent’ for histone marks. Peaks were called against a negative IgG control, generated in each experiment for each condition.

## Supplemental Figure Legends

**Fig. S1:**
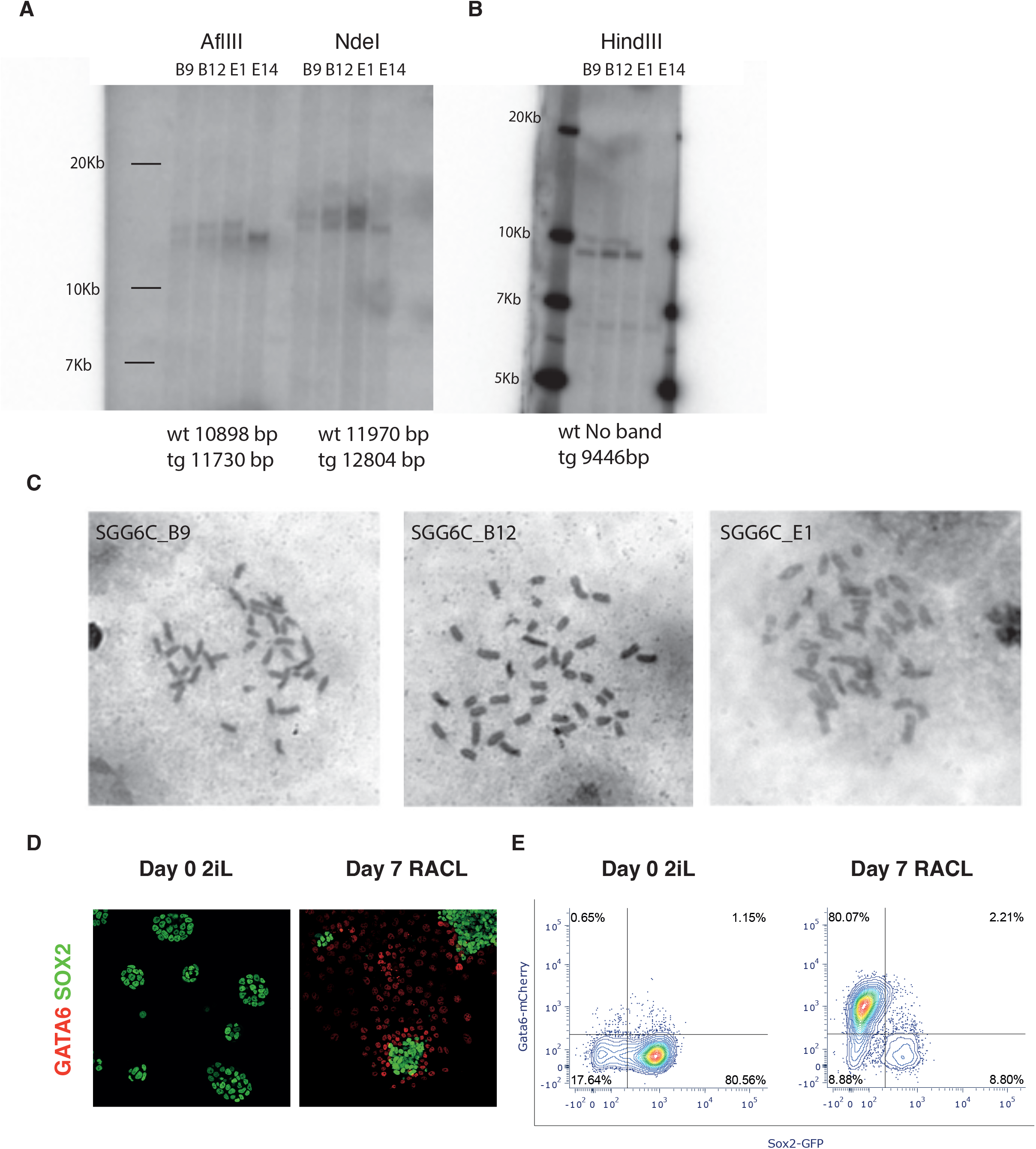
**A-B.** Southern blots for the Gata6::mCherry construct for (A) the external probe and (B) the internal probe, in the cherry sequence resulting in 3 good SGGC clones (B9, B12 & E1). **C.** Bright field images of the correct karyotyping of the 3 SGGC clones. **D.** Representative images of the SGGC lines in 2iLIF (naive ESCs) and RACL conditions (nEnd). **E.** Representative flow cytometry plots of the SGGC cell line in 2iLIF and RACL conditions.

**Fig. S2.**
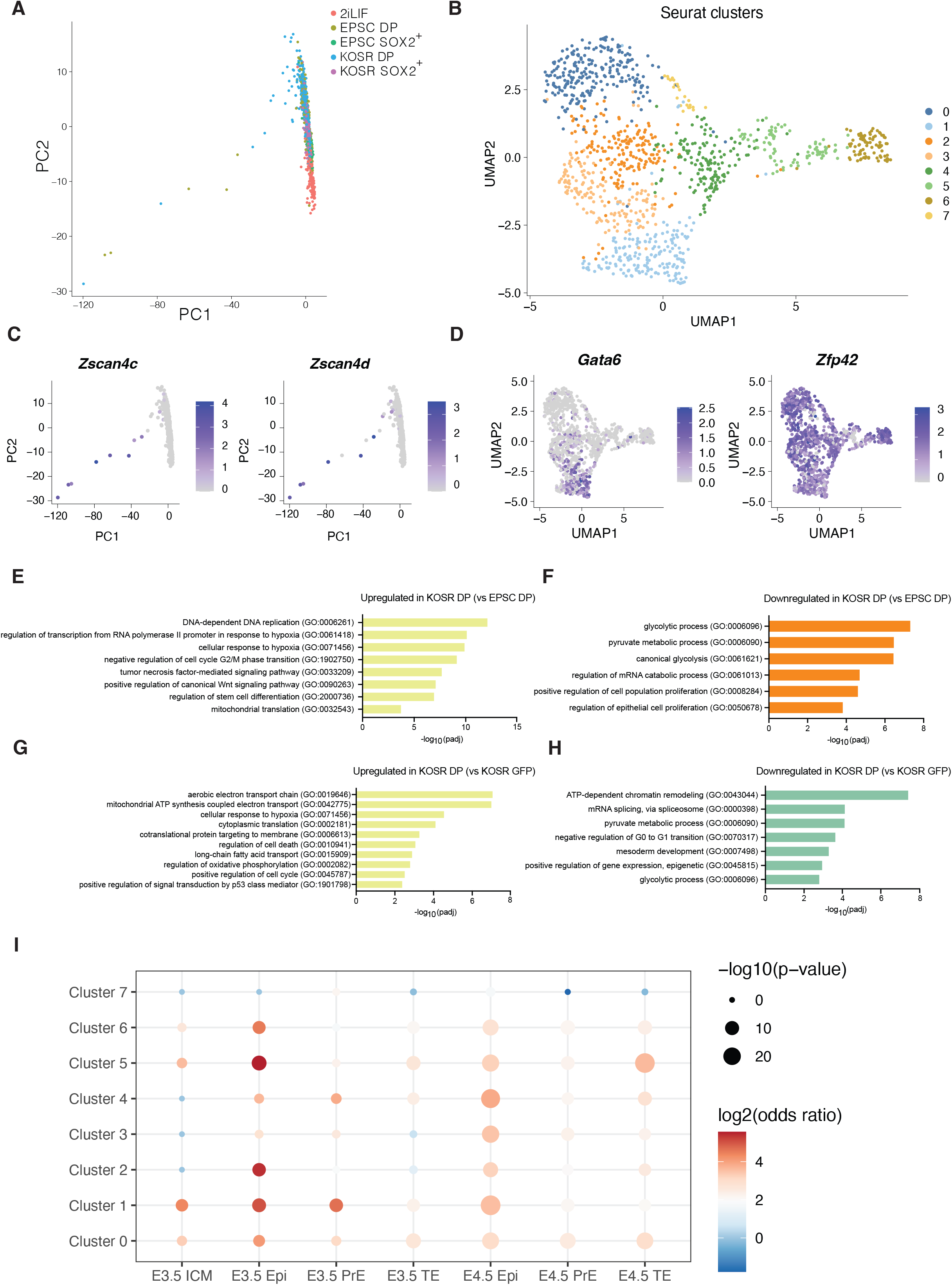
**A.** PCA showing the first and second principal components. PC1 separates 2CLCs from the rest. **B.** UMAP dimensional embedding where colouring represents Louvain clustering. **C.** Expression of 2C genes (*Zscan4c, Zscan4d*) projected on PCA plots. **D.** Expression of selected markers (*Gata6* and *Zfp42*) on UMAP plots. **E-F.** GO term enrichment for KOSR DP cells compared with EPSCM DP cells. **G-H.** GO term enrichment of KOSR DP compared with KOSR SOX2^+^. **I.** Gene overlap analysis comparing unsupervised clusters of our scRNA-seq cluster in B. against differentially expressed genes to *in vivo* scRNA-seq of pre-implantation blastocysts (*Nowotschin et. al 2019*).

**Fig. S3.**
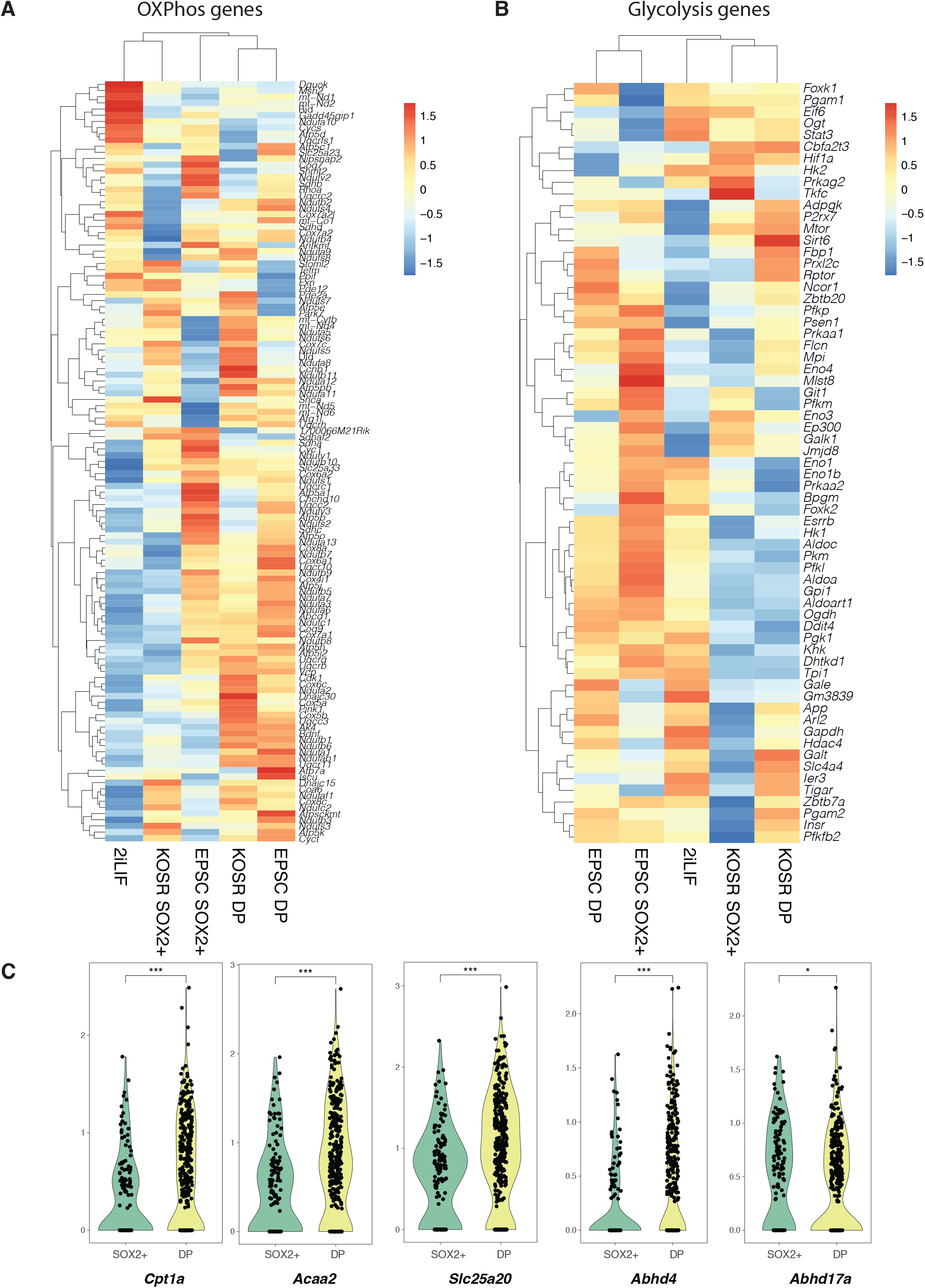
**A.** Heatmap showing oxidative phosphorylation genes expression in the different media conditions. **B.** Heatmap showing glycolysis genes. **C.** Violin plots with different examples of lipid metabolism genes distribution in DP or SOX2^+^ cells in KOSR media. Mann-Whitney U statistical test. * = p-value 0.01, *** = p-value 0.001

**Fig. S4.**
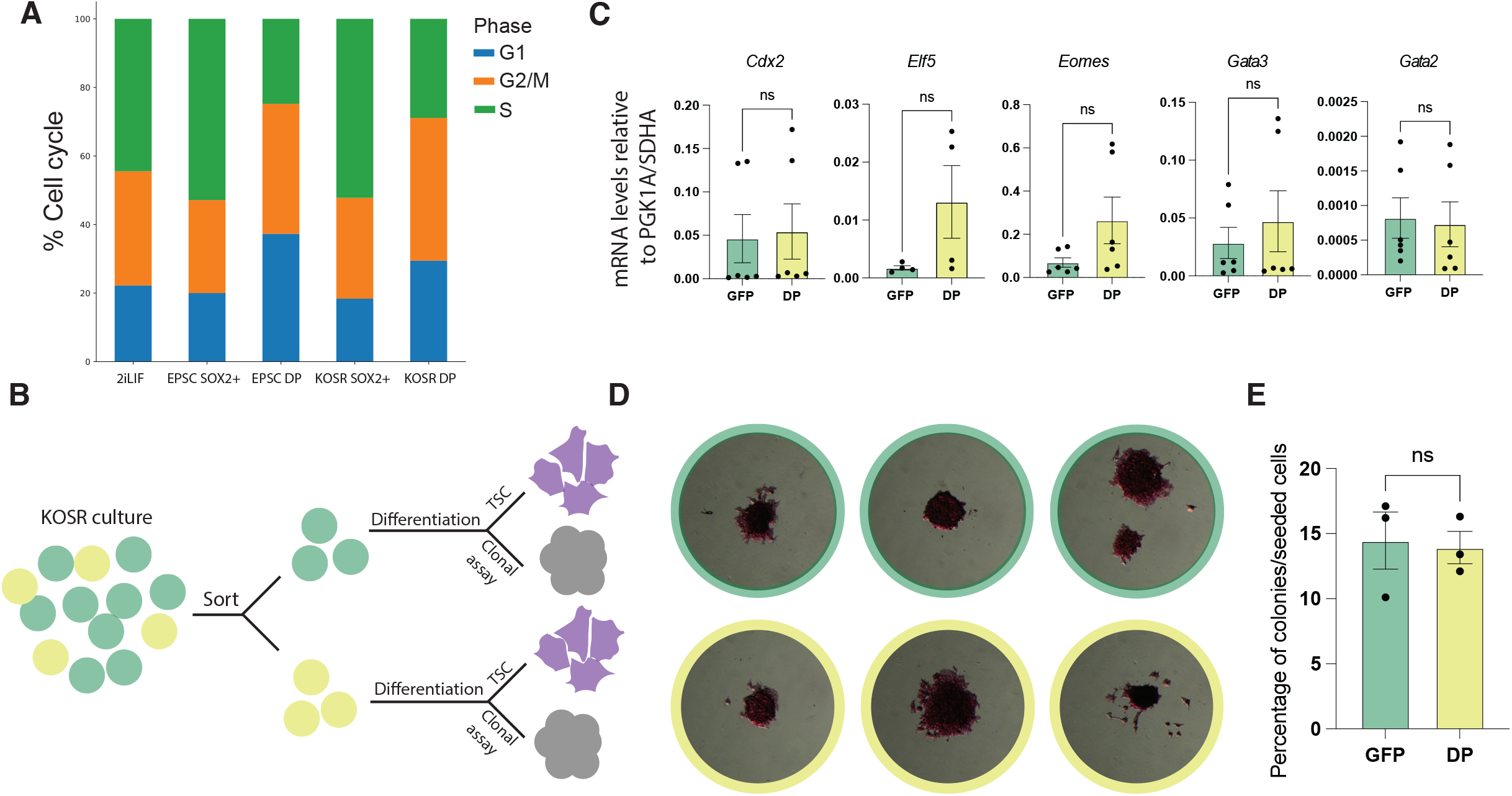
**A**. Distribution of the cell cycle phases in different sorted populations based on sc-RNA-seq data. **B.** Schematic cartoon of the TSC differentiation and clonal assay after sorting KOSR cells by FACS. **C.** Relative mRNA levels of TSC markers of the sorted SOX2^+^ or DP sorted populations after 6 days of TSC differentiation. **D.** Representative images of colonies formed after 1 week of clonal growth after sorting by FACS and AP staining. Colonies that were originally sorted for SOX2^+^ cells have green frames, and colonies coming from DP sorted cells have yellow frames. **E.** Quantification of the AP staining from the clonal assay.

## Tables

**Table 1** shows a list of the closest Epi genes to the cobound peaks for SOX2 and GATA6 in DP cells. It also shows how many SOX2 and GATA6 motifs we find in these regions. Genes colored in yellow are differentially upregulated in DP over SOX2^+^ cells in the scRNA-seq. The ones colored in green are differentially upregulated in SOX2^+^ over DP cells.

**Table 2** shows a list of the closest PrE genes to the cobound peaks for SOX2 and GATA6 in DP cells. It also shows how many SOX2 and GATA6 motifs we find in these regions. Genes colored in yellow are differentially upregulated in DP over SOX2^+^ cells in the scRNA-seq. The ones colored in green are differentially upregulated in SOX2^+^ over DP cells.

**Table 3** compares SOX2 and GATA6 binding peaks from our study with the SOX2 and GATA6 peaks of the GATA6 overexpression time course from *Thompson. et al 2022*.

## References

Anderson, K. G. V., Hamilton, W. B., Roske, F. V., Azad, A., Knudsen, T. E., Canham, M. A., Forrester, L. M. and Brickman, J. M. (2017). Insulin fine-tunes self-renewal pathways governing naive pluripotency and extra-embryonic endoderm. Nat. Cell Biol. 19, 1164–1177.

Ar, Q. and Im, H. (2010). BEDTools: a flexible suite of utilities for comparing genomic features. Bioinforma. Oxf. Engl. 26,.

Boroviak, T., Stirparo, G. G., Dietmann, S., Hernando-Herraez, I., Mohammed, H., Reik, W., Smith, A., Sasaki, E., Nichols, J. and Bertone, P. (2018). Single cell transcriptome analysis of human, marmoset and mouse embryos reveals common and divergent features of preimplantation development. Dev. Camb. Engl. 145, dev167833.

Calo, E. and Wysocka, J. (2013). Modification of Enhancer Chromatin: What, How, and Why? Mol. Cell 49, 825–837.

Canham, M. A., Sharov, A. A., Ko, M. S. H. and Brickman, J. M. (2010). Functional Heterogeneity of Embryonic Stem Cells Revealed through Translational Amplification of an Early Endodermal Transcript. PLoS Biol. 8, e1000379.

Chen, J., Zhang, Z., Li, L., Chen, B.-C., Revyakin, A., Hajj, B., Legant, W., Dahan, M., Lionnet, T., Betzig, E., et al. (2014). Single-Molecule Dynamics of Enhanceosome Assembly in Embryonic Stem Cells. Cell 156, 1274–1285.

Coronado, D., Godet, M., Bourillot, P.-Y., Tapponnier, Y., Bernat, A., Petit, M., Afanassieff, M., Markossian, S., Malashicheva, A., Iacone, R., et al. (2013). A short G1 phase is an intrinsic determinant of naïve embryonic stem cell pluripotency. Stem Cell Res. 10, 118–131.

Dietrich, J.-E. and Hiiragi, T. (2007). Stochastic patterning in the mouse pre-implantation embryo. Dev. Camb. Engl. 134, 4219–31.

Genet, M. and Torres-Padilla, M.-E. (2020). The molecular and cellular features of 2-cell-like cells: a reference guide. Dev. Camb. Engl. 147, dev189688.

Georgiou, G. and van Heeringen, S. J. (2016). fluff: exploratory analysis and visualization of high-throughput sequencing data. PeerJ 4, e2209.

H, L., B, H., A, W., T, F., J, R., N, H., G, M., G, A. and R, D. (2009). The Sequence Alignment/Map format and SAMtools. Bioinforma. Oxf. Engl. 25,.

Hamilton, W. B., Mosesson, Y., Monteiro, R. S., Emdal, K. B., Knudsen, T. E., Francavilla, C., Barkai, N., Olsen, J. V. and Brickman, J. M. (2019). Dynamic lineage priming is driven via direct enhancer regulation by ERK. Nature 575, 355–360.

Hayashi, K., Ohta, H., Kurimoto, K., Aramaki, S. and Saitou, M. (2011). Reconstitution of the Mouse Germ Cell Specification Pathway in Culture by Pluripotent Stem Cells. Cell 146, 519–532.

Heinz, S., Benner, C., Spann, N., Bertolino, E., Lin, Y. C., Laslo, P., Cheng, J. X., Murre, C., Singh, H. and Glass, C. K. (2010). Simple Combinations of Lineage-Determining Transcription Factors Prime cis-Regulatory Elements Required for Macrophage and B Cell Identities. Mol. Cell 38, 576–589.

Janssens, D. and Henikoff, S. (2019). CUT&RUN: Targeted in situ genome-wide profiling with high efficiency for low cell numbers. protocols.io.

Keren-Shaul, H., Kenigsberg, E., Jaitin, D. A., David, E., Paul, F., Tanay, A. and Amit, I. (2019). MARS-seq2.0: an experimental and analytical pipeline for indexed sorting combined with singlecell RNA sequencing. Nat. Protoc. 14, 1841–1862.

Knudsen, T. E. and Brickman, J. M. (2020). Can a Cell Put Its Arms around a Memory? Cell Stem Cell 26, 609–610.

Langmead, B. and Salzberg, S. L. (2012). Fast gapped-read alignment with Bowtie 2. Nat. Methods 9, 357–359.

Leese, H. J. (2012). Metabolism of the preimplantation embryo: 40 years on. Reproduction 143, 417–427.

Li, M. and Belmonte, J. C. I. (2017). Ground rules of the pluripotency gene regulatory network. Nat. Rev. Genet. 18, 180–191.

Linneberg-Agerholm, M., Wong, Y. F., Herrera, J. A. R., Monteiro, R. S. and Anderson, K. G. V. (2019). Naïve human pluripotent stem cells respond to Wnt, Nodal and LIF signalling to produce expandable naïve extra-embryonic endoderm. Development 146,.

Liu, N. (2019). Library Prep for CUT&RUN with NEBNext^®^ Ultra™ II DNA Library Prep Kit for Illumina^®^ (E7645). protocols.io.

Martin, M. (2011). Cutadapt removes adapter sequences from high-throughput sequencing reads. EMBnet.journal 17, 10.

Martin Gonzalez, J., Morgani, S. M., Bone, R. A., Bonderup, K., Abelchian, S., Brakebusch, C. and Brickman, J. M. (2016). Embryonic Stem Cell Culture Conditions Support Distinct States Associated with Different Developmental Stages and Potency. Stem Cell Rep. 7, 177–191.

Meers, M. P., Tenenbaum, D. and Henikoff, S. (2019). Peak calling by Sparse Enrichment Analysis for CUT&RUN chromatin profiling. Epigenetics Chromatin 12, 42.

Morgani, S. M. and Brickman, J. M. (2015). LIF supports primitive endoderm expansion during pre-implantation development. Development 142, 3488–3499.

Morgani, S. M., Canham, M. A., Nichols, J., Sharov, A. A., Migueles, R. P., Ko, M. S. H. and Brickman, J. M. (2013). Totipotent Embryonic Stem Cells Arise in Ground-State Culture Conditions. Cell Rep. 3, 1945–1957.

Morgani, S., Nichols, J. and Hadjantonakis, A.-K. (2017). The many faces of Pluripotency: in vitro adaptations of a continuum of in vivo states. BMC Dev. Biol. 17,.

Nichols, J. and Smith, A. (2011). The origin and identity of embryonic stem cells. Development 138, 3–8.

Nowotschin, S., Setty, M., Kuo, Y.-Y., Liu, V., Garg, V., Sharma, R., Simon, C. S., Saiz, N., Gardner, R., Boutet, S. C., et al. (2019). The emergent landscape of the mouse gut endoderm at singlecell resolution. Nature 1.

Perera, M., Nissen, S. B., Proks, M., Pozzi, S., Monteiro, R. S., Trusina, A. and Brickman, J. M. (2022). Transcriptional Heterogeneity and Cell Cycle Regulation as Central Determinants of Primitive Endoderm Priming. 2022.04.03.486894.

Pietenpol, J. A. and Stewart, Z. A. (2002). Cell cycle checkpoint signaling:: Cell cycle arrest versus apoptosis. Toxicology 181–182, 475–481.

Posfai, E., Schell, J. P., Janiszewski, A., Rovic, I., Murray, A., Bradshaw, B., Yamakawa, T., Pardon, T., El Bakkali, M., Talon, I., et al. (2021). Evaluating totipotency using criteria of increasing stringency. Nat. Cell Biol. 23, 49–60.

Ramírez, F., Ryan, D. P., Grüning, B., Bhardwaj, V., Kilpert, F., Richter, A. S., Heyne, S., Dündar, F. and Manke, T. (2016). deepTools2: a next generation web server for deep-sequencing data analysis. Nucleic Acids Res. 44, W160–165.

Riveiro, A. R. and Brickman, J. M. (2020). From pluripotency to totipotency: an experimentalist’s guide to cellular potency. Development 147,.

Robinson, J. T., Thorvaldsdóttir, H., Winckler, W., Guttman, M., Lander, E. S., Getz, G. and Mesirov, J. P. (2011). Integrative genomics viewer. Nat. Biotechnol. 29, 24–26.

Skene, P. J. and Henikoff, S. (2017). An efficient targeted nuclease strategy for high-resolution mapping of DNA binding sites. eLife 6, e21856.

Southern, E. (2006). Southern blotting. Nat. Protoc. 1, 518–525.

Tanaka, S., Kunath, T., Hadjantonakis, A.-K., Nagy, A. and Rossant, J. (1998). Promotion of Trophoblast Stem Cell Proliferation by FGF4. Science 282, 2072–2075.

Thompson, J. J., Lee, D. J., Mitra, A., Frail, S., Dale, R. K. and Rocha, P. P. (2022). Extensive cobinding and rapid redistribution of NANOG and GATA6 during emergence of divergent lineages. Nat. Commun. 13,.

White, M. D., Angiolini, J. F., Alvarez, Y. D., Kaur, G., Zhao, Z. W., Mocskos, E., Bruno, L., Bissiere, S., Levi, V. and Plachta, N. (2016). Long-Lived Binding of Sox2 to DNA Predicts Cell Fate in the Four-Cell Mouse Embryo. Cell 165, 75–87.

Yang, J., Ryan, D. J., Wang, W., Tsang, J. C. H., Lan, G., Masaki, H., Gao, X., Antunes, L., Yu, Y., Zhu, Z., et al. (2017a). Establishment of mouse expanded potential stem cells. Nature 550,.

Yang, Y., Liu, B., Xu, J., Wang, J., Wu, J., Shi, C., Xu, Y., Dong, J., Wang, C., Lai, W., et al. (2017b). Derivation of Pluripotent Stem Cells with In Vivo Embryonic and Extraembryonic Potency. Cell 169, 243–257.e25.

Ying, Q. L., Wray, J., Nichols, J., Batlle-Morera, L., Doble, B., Woodgett, J., Cohen, P. and Smith, A. (2008). The ground state of embryonic stem cell self-renewal. Nature 453, 519–523.

